# Machine learning classifies subcellular Golgi morphology from imaging flow cytometry during inflammatory activation: an exploratory study

**DOI:** 10.64898/2026.10.01.756023

**Authors:** Xu Zhang, Mengchen Fan, Baocheng Geng, Pulin Che

**Affiliations:** Department of Anesthesiology and Perioperative Medicine, Division of Molecular and Translational Biomedicine, Heersink School of Medicine, University of Alabama at Birmingham, Birmingham, AL, USA; Department of Computer Science, University of Alabama at Birmingham, Birmingham, AL, USA

**Keywords:** Golgi apparatus, imaging flow cytometry, deep learning, inflammatory activation, organelle morphology, cross-species transfer learning, immune cell phenotyping

## Abstract

Inflammatory activation remodels the Golgi apparatus to meet increased secretory demand, yet whether these structural changes carry computationally accessible information about immune cell state has not been explored. Here we show that machine learning applied to imaging flow cytometry images of the Golgi can distinguish lipopolysaccharide (LPS) treated from control cells. We trained models on Golgi images from human THP-1 monocytes before and 16 h after LPS stimulation in vitro, and from circulating myeloid cells of mice 24 h after intratracheal LPS or saline in vivo. In each dataset, the best-performing model was a deep learning model trained directly on images rather than a classifier using predefined morphometric features. Cross-domain performance varied with the model, transfer direction and use of labelled target data: several deep learning models trained on mouse images classified THP-1 cells above chance without changing model parameters. Grad-CAM highlighted regions within or adjacent to the GM130 fluorescence pattern. These findings indicate that GM130 images carry information associated with LPS treatment that supports classification within and across the two datasets. To our knowledge, this is the first application of supervised deep learning to imaging flow cytometry images of a subcellular organelle for inflammatory phenotyping, establishing organelle level image analysis as a potential complement to molecular assays for immune cell classification.

## Introduction

The Golgi apparatus sorts and dispatches secreted proteins as they move from the endoplasmic reticulum (ER) to the plasma membrane ^1–3^. Although this organelle was once regarded as structurally fixed, it is now known to remodel as secretory demand changes, with the ribbon expanding, fragmenting and reassembling ^4–6^. Innate immune activation is a setting where secretory demand rises sharply. In macrophages and monocytes, Toll-like receptor 4 detects lipopolysaccharide (LPS) and shifts the cell toward cytokine production ^7, 8^. Newly made cytokines pass through the Golgi, and the organelle remodels as traffic increases. This remodeling has been documented in several systems, but predominantly through qualitative assessment; whether computational methods can systematically distinguish an activated cell from a resting one on the basis of organelle morphology remains untested. If these structural changes are consistent, they constitute computationally accessible information about cell state. We therefore asked whether machine learning applied to Golgi images could distinguish activated from resting immune cells, establishing organelle-level morphology-based phenotyping as a complement to established molecular assays.

Answering this question requires an imaging platform that preserves subcellular spatial information while maintaining sufficient throughput for statistical analysis. Conventional flow cytometry reduces each cell’s fluorescence to a single intensity value, discarding where in the cell the signal originated ^9^. Imaging flow cytometry records an image of every cell instead, preserving subcellular spatial information at comparable throughput ^10, 11^. Platforms such as the ImageStreamX combine the throughput of flow cytometry with the spatial detail of microscopy, producing morphometric features from thousands of cells per sample ^9, 12, 13^. Golgi morphometry using the cis-Golgi matrix protein GM130 is established on this platform ^14^. Deep learning applied to ImageStreamX data has classified cell cycle phase with high accuracy ^15^, and conventional machine learning has classified white blood cell subtypes from label-free ImageStreamX images ^16^, showing that this platform supports image-based classification.

In subcellular organelle analysis, unsupervised machine learning applied to confocal images of a single Golgi marker revealed ten distinct phenotypic states ^17^. Deep learning models have discriminated cell cycle state, with gradient-weighted class activation mapping (Grad-CAM) used to interpret their predictions ^18^. A protocol for deep learning based classification of Golgi spatial distribution changes has been described for confocal microscopy images ^19^. Machine learning applied to whole cell morphology classifies macrophage activation at single cell resolution, with detection that agrees with cytokine secretion ^20^, and label-free imaging flow cytometry data support macrophage phenotype classification ^21^. These studies suggest that machine learning can differentiate cellular functional states from images and that imaging flow cytometry provides suitable throughput for such analyses. However, no study has applied machine learning to imaging flow cytometry images of a subcellular organelle to classify inflammatory activation.

In this study, using GM130 labelled Golgi images acquired by imaging flow cytometry, we explore the potential of machine learning to classify subcellular organelle morphology changes associated with inflammatory activation. We first examine whether an imaging and classification pipeline can capture and distinguish activation states in a controlled *in vitro* system using the THP-1 cell line, where both the stimulus and the cell population are defined. We then test whether the same methodological approach extends to primary cells from a living animal, where the stimulus is systemic and the cell population is heterogeneous. As an exploratory proof of concept, this study shows that in both systems Golgi images contain patterns that help distinguish activated immune cells from control, and examines whether learned image representations transfer across species and experimental contexts.

## Results

### An imaging flow cytometry and machine learning pipeline classifies GM130 morphology in LPS activated THP-1 monocytes

We first asked whether imaging flow cytometry could provide Golgi images at sufficient quality and throughput to classify inflammatory activation in individual cells using deep learning. The analytical pipeline, from sample acquisition through model training and evaluation, is summarized in **Fig. 1A**. THP- 1 monocytes were collected, fixed, and stained at 0 and 16 hours after LPS stimulation, followed by acquisition on an ImageStreamX Mark II at 60× magnification. Representative single-cell images confirmed that the GM130 fluorescence signal was preserved at a resolution sufficient to resolve subcellular morphology, alongside nuclear (DAPI), brightfield and viability channels **(Fig. 1B)**. Focused, in-frame single-cell events were then selected through a prespecified quality-control gating workflow. To visualize patterns associated with treatment, we used a supervised feature encoder to summarize how each cell’s IDEAS measurements contributed to its predicted class. UMAP and t-SNE projections of these encoded features showed separation between the 0 h and 16 h conditions **(Fig. 1C).**

**Figure 1.**
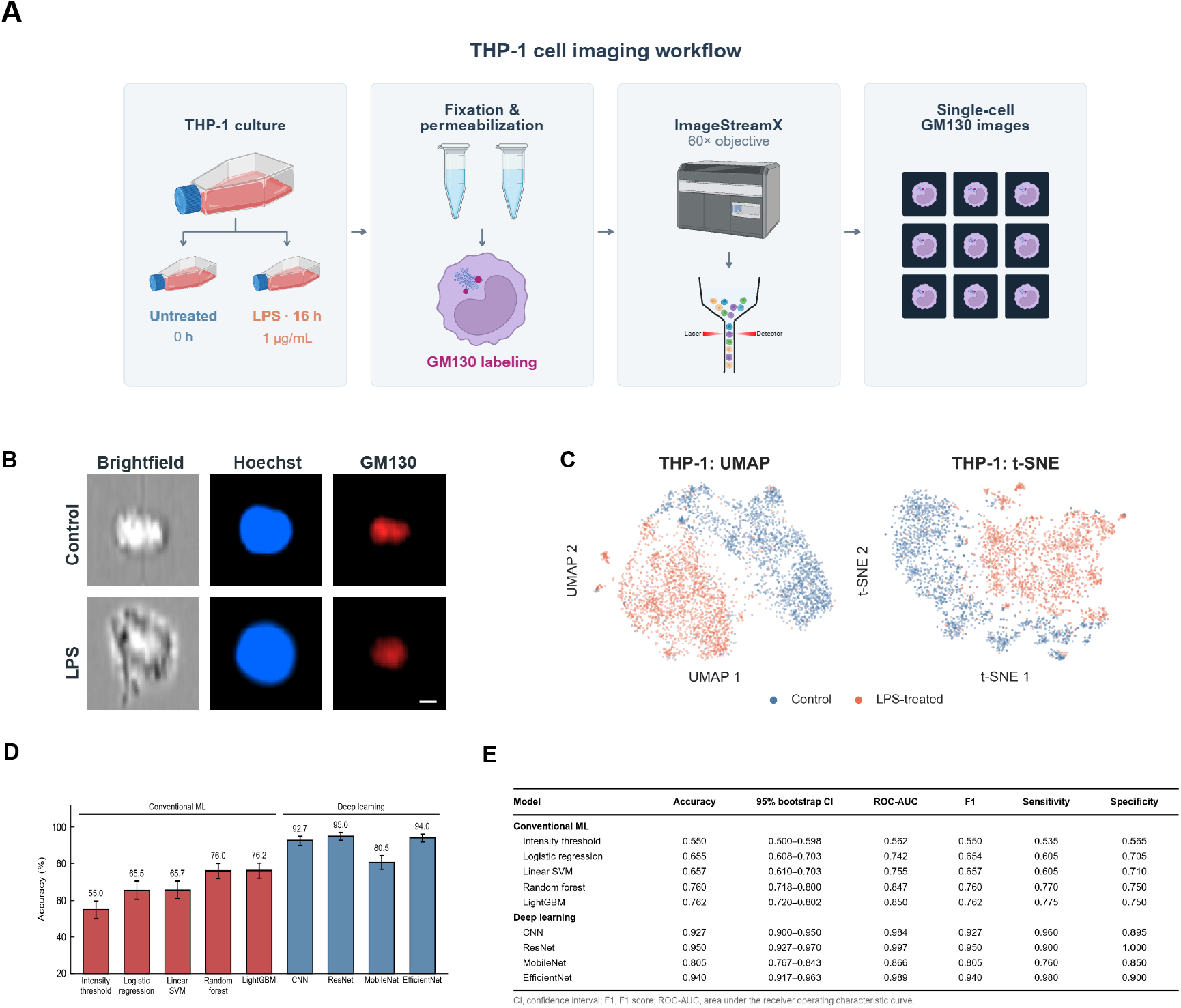
Imaging flow cytometry and machine learning classification of GM130 morphology in LPS-treated THP-1 monocytes. **A**, Experimental workflow: THP-1 monocytes were left untreated (0 h) or exposed to LPS (1 µg/mL, 16 h), fixed, immunolabelled for GM130 and imaged on the ImageStreamX at 60× magnification; GM130 images of individual cells were used for classification. **B**, Representative single-cell images from the 0 h and 16 h conditions, showing the GM130-AF594, Hoechst, brightfield and Ghost Dye Red 780 channels. **C**, UMAP and t-SNE projections of encoded IDEAS features, coloured by condition; these supervised projections are exploratory. **D**, Test accuracy of five conventional classifiers trained on IDEAS features and four deep learning models trained on GM130 images, evaluated on the held-out test set (400 images, 200 per condition); error bars, 95% bootstrap confidence intervals from 2,000 stratified resamples. **E**, Accuracy, 95% bootstrap CI, ROC-AUC, F1 score, sensitivity and specificity of the same models on the same test set.

We selected 4,000 images and divided them into training, validation and held-out test sets. Deep learning models trained directly on the GM130 channel outperformed conventional classifiers trained on morphometric features extracted by the ImageStreamX IDEAS software. ResNet-18, selected using the validation set, achieved a test accuracy of 0.950 and a ROC-AUC of 0.997, whereas the strongest conventional classifier, LightGBM, reached an accuracy of 0.762 **(Figs. 1D, E)**. These results show that GM130 images contain information that distinguishes LPS-treated from control THP-1 cells, with deep learning applied directly to images outperforming classifiers trained on the IDEAS features in this dataset.

### A single intratracheal LPS challenge produces measurable systemic and pulmonary responses

Having established that the pipeline classified activation state in a defined cell line, we next sought to apply the same approach to primary circulating myeloid cells from an *in vivo* inflammatory model. We chose the intratracheal LPS challenge, a well-established murine model of acute pulmonary inflammation ^22–24^, and verified that it produced the expected inflammatory response under our experimental conditions. Mice received a single intratracheal dose of LPS (1.5 mg/kg) or sterile saline, followed by terminal blood and lung collection at 24 h **(Fig. 2A)**. LPS-treated animals lost body weight over the 24 h challenge period relative to saline controls **(Fig. 2B)**, consistent with the systemic inflammatory response expected from intratracheal endotoxin exposure. Serum levels of the pro-inflammatory cytokines IL-6 and TNF-α, assayed from terminal blood samples, were elevated in the LPS group **(Fig. 2C)**, indicating circulating inflammatory activation. To assess pulmonary involvement, excised lungs were weighed before and after desiccation; the resulting wet-minus-dry mass, a proxy for pulmonary fluid accumulation that does not control differences in lobe size, was elevated in LPS-treated animals **(Fig. 2D)**, suggesting pulmonary edema. Together, these independent measures of weight loss, circulating inflammatory markers and pulmonary fluid accumulation confirmed that the intratracheal LPS model produced a systemic and pulmonary inflammatory response within the 24 h window, validating the use of these samples for subsequent imaging and classification analyses.

**Figure 2.**
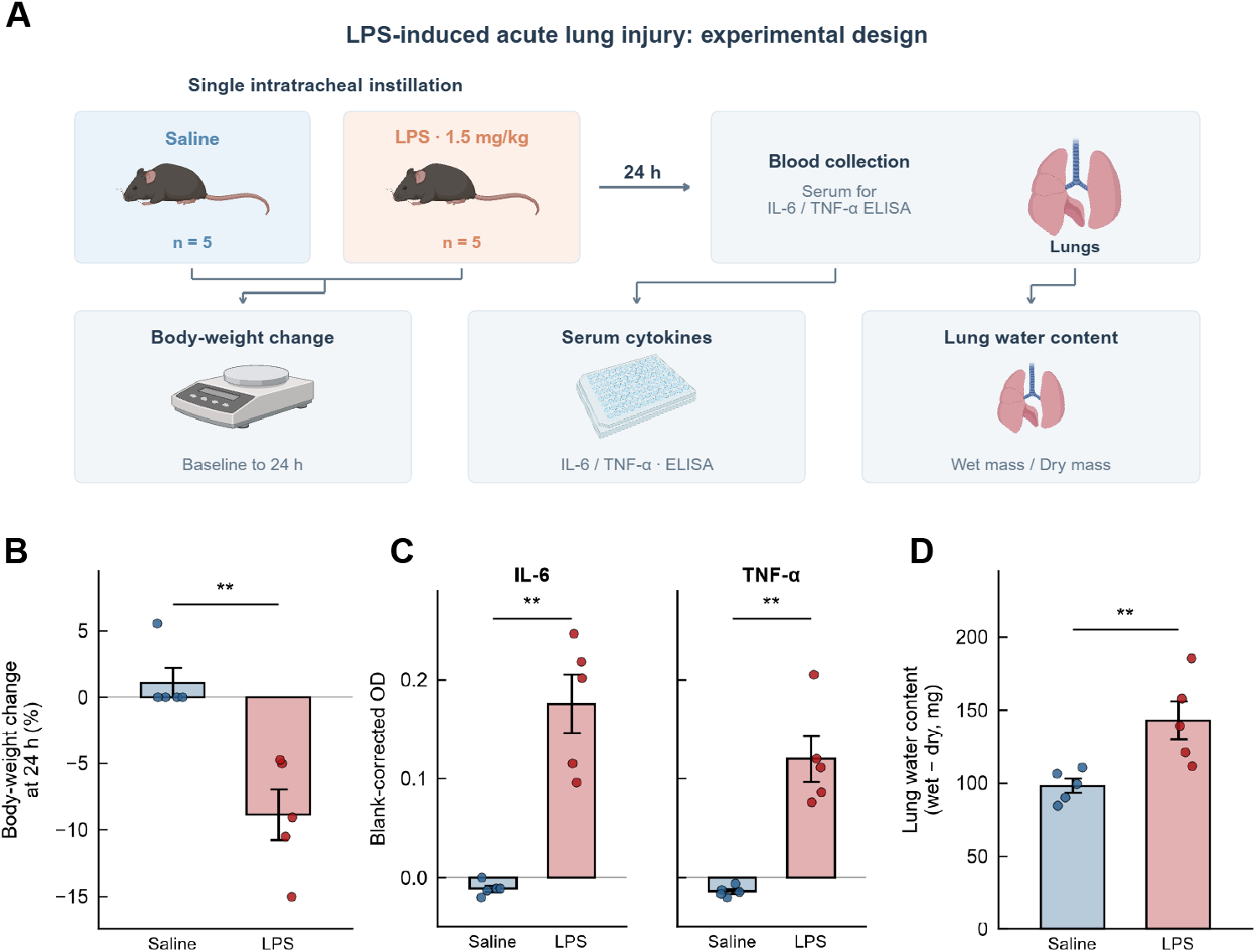
Validation of the intratracheal LPS mouse model. **A**, Experimental design: mice received a single intratracheal instillation of LPS (1.5 mg/kg) or sterile saline at 0 h, with terminal collection of blood and lungs at 24 h. **B**, Body-weight change from baseline to 24 h. **C**, Serum IL-6 and TNF-α by ELISA, reported as blank-corrected optical density averaged over three technical wells per mouse. D, Lung water content, the difference between the wet and dry mass of the excised lung. In **B-D**, each point is one mouse, bars show the mean and error bars; n = 5 mice per group; **P < 0.01, two-sided Mann-Whitney U test.

### The classification methodology extends to primary circulating myeloid cells from the *in vivo* LPS challenge

We next applied the imaging and classification pipeline to circulating myeloid cells to determine whether the approach could extend to primary cells in an *in vivo* inflammatory context. Peripheral blood collected at 24 h was stained for surface markers and GM130 and acquired on the ImageStreamX **(Fig. 3A)**. Circulating CD45⁺CD11b⁺Ly6G⁻ myeloid cells were identified through a prespecified sequential gating strategy that selected focused single cells, excluded non-viable events, and gated CD45⁺, CD11b⁺ and Ly6G⁻ populations in turn **(Fig. 3B)**, and representative single-cell images confirmed that the GM130 fluorescence signal was preserved alongside the surface-marker, nuclear, brightfield and viability channels in these primary cells **(Fig. 3C)**. We compared two approaches: conventional classifiers using IDEAS morphometric features and deep learning models using GM130 images **(Fig. 3D)**. UMAP and t-SNE projections of encoded IDEAS features showed structure associated with treatment group, with partial overlap between groups **(Fig. 3E)**. The analysis set comprised 1,366 images (371 saline, 995 LPS-treated), split proportionally into training, validation and held-out test sets. EfficientNet achieved an accuracy of 0.885 (95% CI 0.826-0.943) on the held-out test set, whereas conventional classifiers ranged from 0.500 to 0.609. The linear SVM assigned every cell to the LPS class and therefore had an accuracy of 0.500, equal to the majority-only baseline **(Fig. 3F)**. By ROC-AUC, the four deep learning models reached 0.654-0.947, with EfficientNet highest at 0.947, and the conventional classifiers ranged from 0.560 to 0.647 **(Fig. 3G, H)**. These results demonstrated the pipeline classified circulating myeloid cells from LPS-treated and saline-control mice, with deep learning models outperforming conventional classifiers by ROC-AUC.

**Figure 3.**
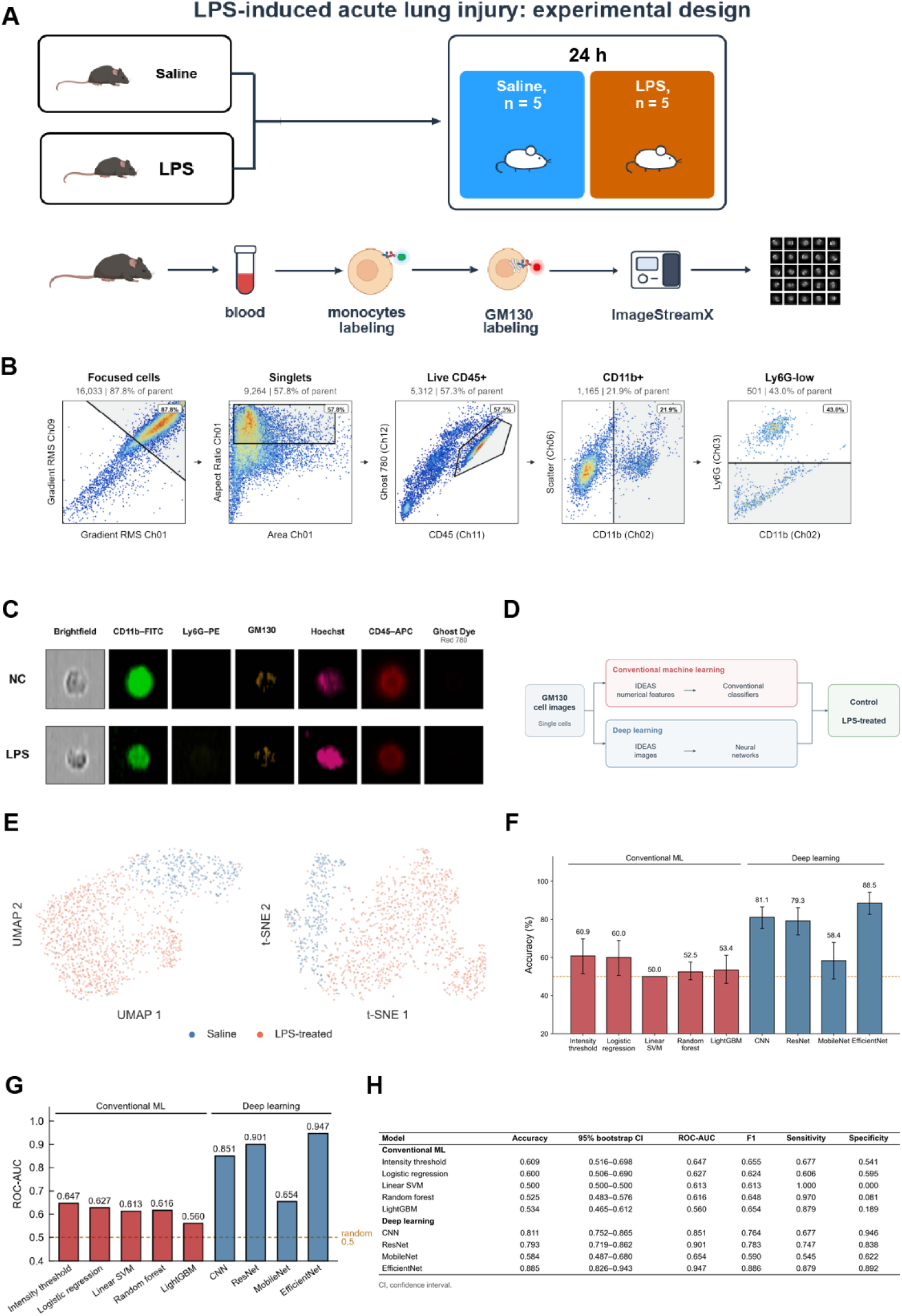
Classification of GM130 morphology in circulating myeloid cells from LPS-challenged mice. **A**, Experimental design and blood-processing workflow: peripheral blood collected 24 h after intratracheal LPS or saline was stained for surface markers and GM130 and acquired on the ImageStreamX. **B**, Sequential gating of CD45⁺CD11b⁺Ly6G⁻ myeloid cells in one representative sample: focused cells, singlets, live CD45⁺ cells, CD11b⁺ cells and Ly6G⁻ cells; percentages are of the parent gate. **C**, Representative single-cell images from the saline and LPS groups, showing the brightfield, CD11b-FITC, Ly6G-PE, GM130, Hoechst, CD45-APC and Ghost Dye Red 780 channels. **D**, Machine learning workflow: conventional classifiers trained on IDEAS morphometric features and deep learning models trained on the GM130 images, both classifying each cell as control or LPS-treated. **E**, UMAP and t-SNE projections of encoded IDEAS features for all 1,366 mouse cells, coloured by group; these supervised projections are exploratory. **F**, Test accuracy of conventional and deep learning models on the held-out mouse test set; error bars, 95% bootstrap confidence intervals from 2,000 stratified resamples; dashed line, majority-only accuracy. **G**, ROC-AUC of the same models; dashed line, 0.5. **H**, Full test metrics.

### Feature importance analysis and conventional microscopy identify condition associated morphometric structure

To understand what image features supported the classification, we assessed which numerical features were most important for prediction. Permutation importance analysis, which quantified the decrease in accuracy after individual features were shuffled, identified minimum spot area and bright detail intensity as the two most influential features in the THP-1 dataset, followed by compactness, gradient RMS and modulation **(Fig. 4A)**. A radar plot summarized the relative importance of the five most important THP-1 features **(Fig. 4B)**. To complement the imaging flow cytometry findings with an independent imaging modality, we examined the same GM130 morphometric features in THP-1 cells processed by conventional immunofluorescence microscopy. Representative images show Golgi structure in untreated and 16 h LPS-treated cells **(Fig. 4C)**. Minimum spot area, the highest-ranked feature in the imaging flow cytometry classifier, was lower after LPS treatment (P = 0.0022, two-sided Wilcoxon rank-sum test, Benjamini-Hochberg q = 0.022 across ten features), and gradient RMS was higher (P = 0.0046, q = 0.023); gradient RMS also showed significant heterogeneity among the three 0 h acquisition batches. Compactness and bright detail intensity did not differ after correction (Fig. 4D). The feature ranked most influential in the imaging flow cytometry classifier corresponds to a change in Golgi organization that is also measurable by conventional fluorescence microscopy.

**Figure 4.**
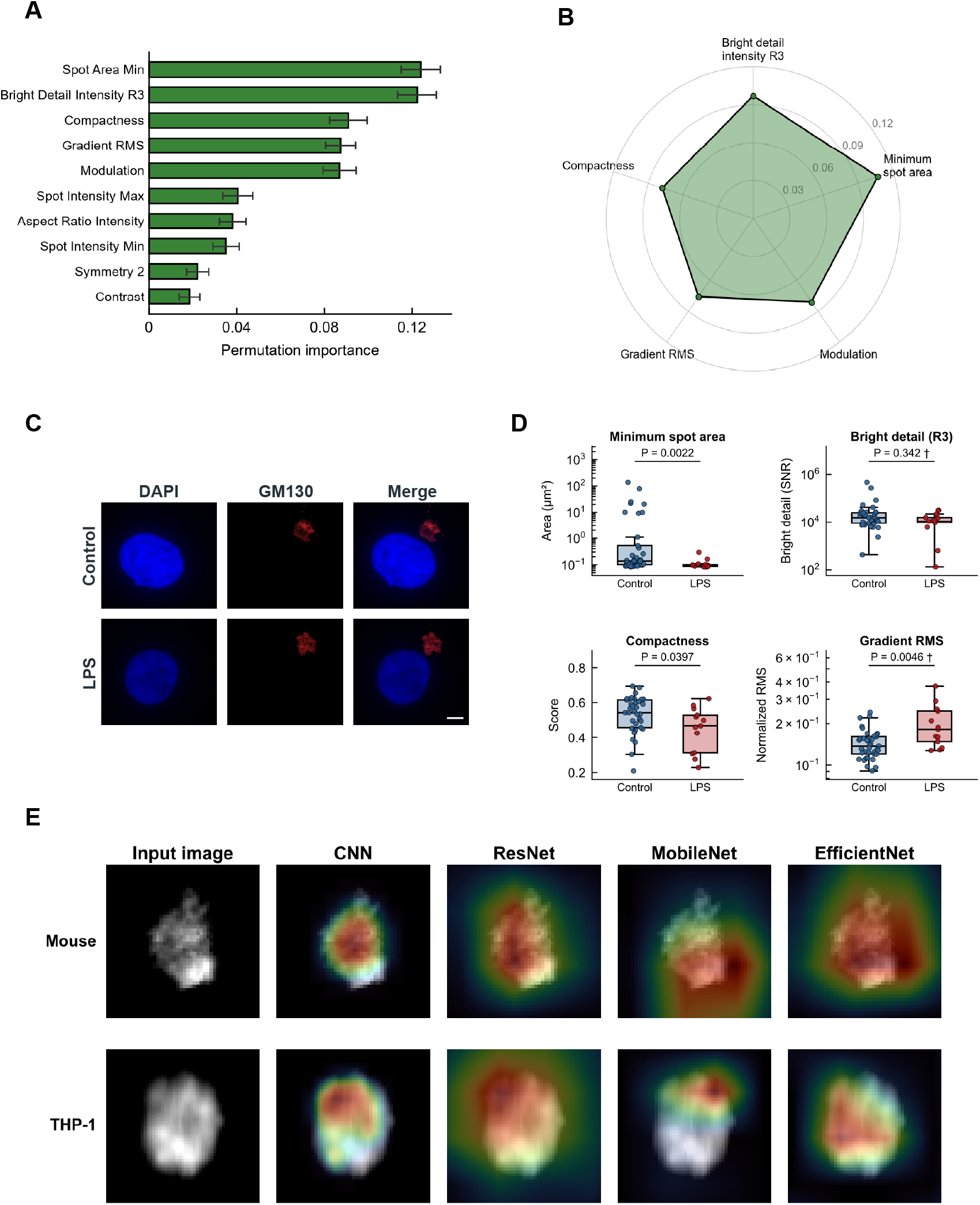
Feature importance, conventional microscopy and Grad-CAM interpretation of the THP-1 classifier. **A**, Permutation importance of the ten IDEAS features for a support vector machine trained on those features, measured as the decrease in balanced accuracy on the held-out THP-1 test set when each feature is shuffled 20 times; bars show the mean and error bars the s.d. **B**, Radar plot of the five most important features from A. **C**, Conventional immunofluorescence images of THP-1 cells at 0 h and 16 h after LPS, stained for DAPI and GM130 (Alexa Fluor 594) and acquired on an ECHO fluorescence microscope at 60× magnification; scale bar, 5 µm. **D**, Field-level medians of the four top-ranked GM130 features at 0 h (n = 37 fields pooled from three acquisition batches) and 16 h (n = 13 fields); each circle is one field, boxes show the median and interquartile range and whiskers extend to 1.5× the interquartile range. P values are from two-sided Wilcoxon rank-sum tests (unadjusted); †, significant heterogeneity among the 0 h acquisition batches (Kruskal-Wallis, Benjamini-Hochberg q < 0.05). Exposure settings were not held constant between conditions, which limits the comparability of intensity-dependent features. **E**, Grad-CAM maps for one representative LPS-treated THP-1 cell and one circulating myeloid cell from an LPS-treated mouse, each classified correctly by all four models, showing the input GM130 image and the maps from the CNN, ResNet, MobileNet and EfficientNet; warm colors mark regions contributing positively to the predicted class (Fig. 4E).

To extend the analysis beyond predefined features, we applied Grad-CAM to visualize the image regions that contributed to the deep learning predictions. In the representative LPS-treated THP-1 and mouse cells, all four architectures (CNN, ResNet, MobileNetV3 and EfficientNet) produced activation maps that localized to regions within or adjacent to the GM130 fluorescence pattern **(Fig. 4E)**, suggesting that these regions contributed to the predictions; the maps do not rule out contributions from fluorescence intensity, background or the morphology of the whole cell. The location and extent of the highlighted regions varied across architectures: the CNN produced compact focal hotspots, whereas EfficientNet highlighted broader areas of the Golgi signal. This architectural variation is consistent with reports that different CNN architectures encode the same image through complementary learned representations ^25^, and illustrates differences in how the models localized image signals associated with the predicted class.

### Machine learning models for cross-domain prediction

To test whether a model trained on one dataset could help classify cells in the other, we compared no, partial and full adaptation between the THP-1 and mouse datasets **(Fig. 5A)**. The source dataset was used for initial training and the target dataset for evaluation. No adaptation kept the source model fixed; partial and full adaptation used labelled target training cells for recalibration or retraining. All three strategies used target validation data to select decision thresholds. Accuracy depended on the model, transfer direction and adaptation strategy **(Fig. 5B; Supplementary Table S1)**. When models trained on mouse images were applied to THP-1 cells, CNN accuracy was 90.2% with fixed parameters and 92.5% after full adaptation, while ResNet improved from 79.5% to 94.8%. MobileNet achieved an accuracy of 92.8% after partial adaptation and 71.0% after full adaptation. When models trained on THP-1 images were applied to mouse cells, the highest accuracy was 82.6% for EfficientNet after partial adaptation, compared with a majority- only baseline of 50.0%. On the independent target test sets, ROC-AUC after full adaptation was higher for all four deep learning models when predicting THP-1 cells using models initially trained on mouse images (0.775-0.993) than in the reverse direction (0.709-0.873) **(Fig. 5C)**. UMAP projections of encoded IDEAS features showed partial separation between saline and LPS-treated mouse cells and clearer separation between control and LPS-treated THP-1 cells **(Fig. 5D)**. For the representative mouse cell evaluated by models trained on THP-1 images, Grad-CAM maps shifted from the peripheral background towards the GM130 signal after adaptation. For the representative THP-1 cell evaluated by models trained on mouse images, the maps already highlighted the cellular signal before adaptation **(Fig. 5E)**. Transfer was asymmetric: models initially trained on mouse images classified THP-1 cells more accurately than the reverse, and adaptation improved performance in both directions.

**Figure 5.**
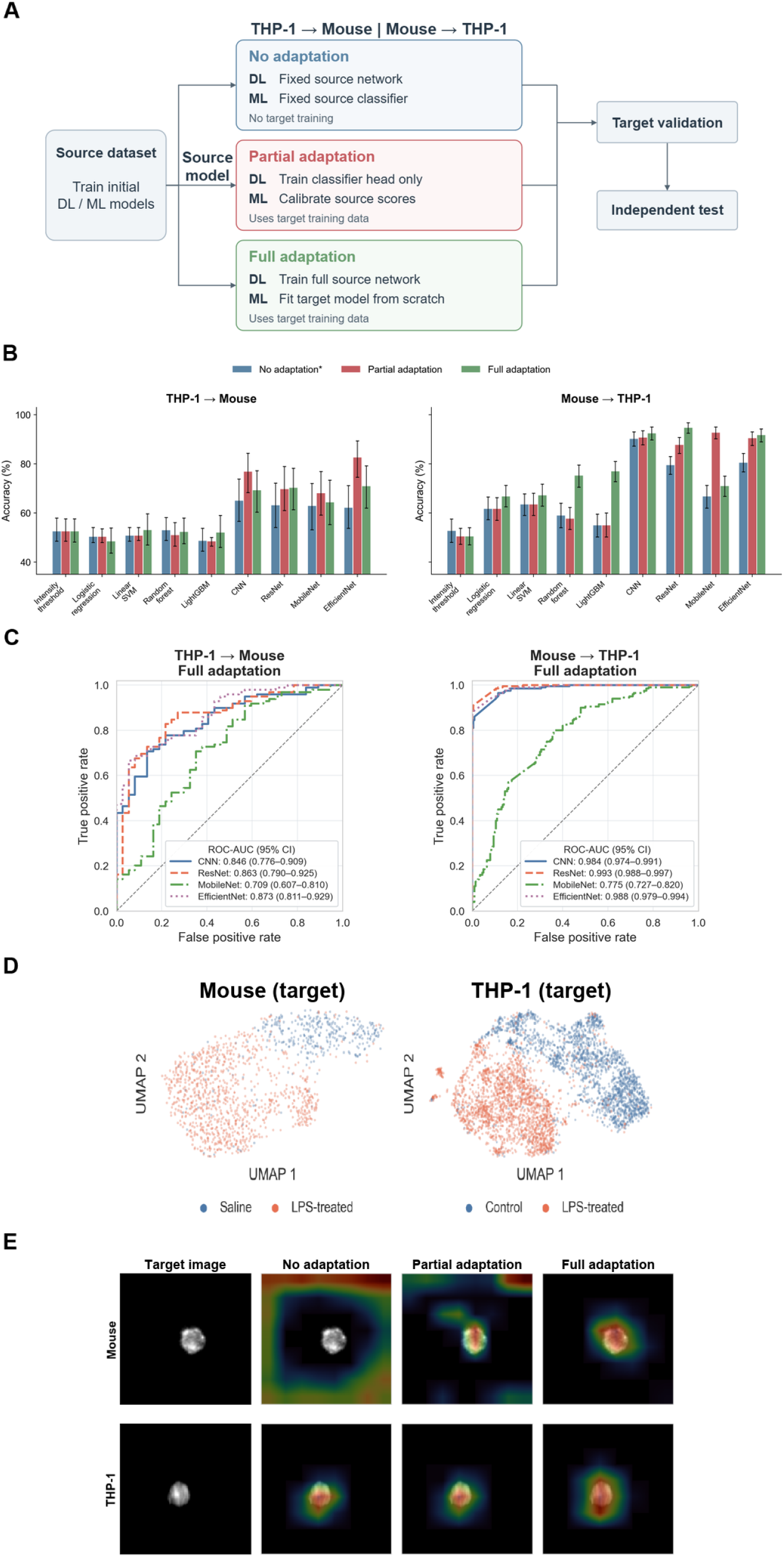
Cross-domain classification between THP-1 cells and mouse myeloid cells. **A**, Bidirectional classification between the THP-1 and mouse datasets under no, partial or full adaptation. The source dataset provides the initial model; the target dataset is the other dataset. No adaptation keeps model parameters fixed; partial adaptation retrains the deep learning classifier head or recalibrates conventional ML prediction scores; full adaptation retrains the entire network or refits the conventional classifier from scratch on target training data. Target validation data were used to select the decision threshold under all three strategies, as marked by the asterisk in B. **B**, Cross-domain test accuracy; error bars, 95% bootstrap confidence intervals from 1,000 stratified resamples of the test set. The majority-only accuracy is 50% for both target test sets. Values are listed in Supplementary Table S1. **C**, ROC curves after full adaptation on independent target test sets (136 mouse cells and 400 THP-1 cells); ROC-AUC confidence intervals use 2,000 stratified resamples. **D**, Separate UMAP projections of encoded IDEAS features, using an encoder fitted on the corresponding target training set. All target cells are displayed (1,366 mouse cells and 4,000 THP-1 cells), including training, validation and test splits; colours indicate treatment group. **E**, ResNet Grad-CAM maps for one representative target cell per direction: a mouse cell evaluated by models trained on THP-1 images (top) and a THP-1 cell evaluated by models trained on mouse images (bottom). The input image and maps under no, partial and full adaptation are shown; warm colours indicate positive contributions to the LPS class score.

## Discussion

In this study, we show that deep learning classifiers trained on imaging flow cytometry images of a single Golgi marker can distinguish inflammatory state in individual immune cells and that these classifiers partially transfer between human and mouse datasets. Prior studies have applied machine learning to the morphology of whole cells for macrophage classification ^20, 21^ and used unsupervised methods to resolve Golgi phenotypic states from confocal microscopy ^17^. To our knowledge, this work introduces two elements not previously reported: the application of deep learning to imaging flow cytometry images of a subcellular organelle marker for inflammatory phenotyping, and an assessment of whether the resulting image features transfer across species. The central finding is that images of a Golgi marker contain information associated with LPS treatment that is useful for cell classification, with some of this information shared between the THP-1 and mouse datasets.

What the models capture can be partly inferred from the classifier comparison and the underlying biology. The best-performing model in each dataset was a deep learning model (ResNet in THP-1, EfficientNet in mouse), suggesting that the GM130 images contain useful information not fully captured by the selected morphometric features. This information may include spatial patterns, texture and fluorescence intensity. The biological basis for this readout is grounded in the established reorganization of the secretory apparatus during innate immune activation. LPS-driven cytokine production requires Golgi remodeling to accommodate increased trafficking ^26–28^, and inflammatory activation restructures interconnected organelle networks ^29^. The reduction in minimum area of GM130-positive structures detected here is compatible with this secretory remodeling, and its independent measurement by conventional microscopy (Fig. 4C, D) supports the interpretation that the classifiers drew on structural changes rather than acquisition artifacts alone.

Conserved Golgi remodeling during inflammation may also explain why some image features transferred between the THP-1 and mouse datasets. Several deep learning models trained on mouse images classified THP-1 cells above chance with model parameters held fixed, suggesting that the two datasets share image patterns associated with LPS treatment. Adaptation further improved performance for some models, although the gains varied across architectures and transfer directions. After full adaptation, ROC-AUC was higher when applying models trained on mouse images to THP-1 cells than in the reverse direction. UMAP shows patterns associated with treatment within each target dataset, while Grad-CAM highlights image regions contributing to predictions in selected cells (Fig. 5D, E).

This work has several limitations. Cells were split at the image level rather than by animal or acquisition batch, so the results represent within-experiment classification, not generalization to new animals or batches. The models may draw on fluorescence intensity, background or acquisition conditions alongside Golgi morphology, and the comparison does not establish that subcellular imaging outperforms whole-cell imaging. In the cross-domain experiments, adapted results do not establish an advantage of source pre-training over training on the target data alone. Single time points and one organelle marker further bound these conclusions; animal-grouped validation and independent replication are needed.

In summary, deep learning classifiers distinguished LPS-treated from control cells using imaging flow cytometry images of a single Golgi marker in both THP-1 and mouse datasets. Deep learning outperformed conventional classifiers trained on predefined morphometric features, and the top-ranked feature was independently confirmed by conventional fluorescence microscopy. Cross-domain experiments showed that some of this learned information transferred between species, with several models classifying THP-1 cells above chance using parameters learned from mouse images alone. These results demonstrate that Golgi morphology carries single-cell information about inflammatory activation accessible to machine learning. Extending this approach to additional markers, time points and independent cohorts would test whether the principle generalizes beyond the conditions studied here.

## Materials and Methods

### Cell culture and LPS treatment

Human THP-1 monocytic cells were maintained in suspension in RPMI 1640 supplemented with 10% heat-inactivated fetal bovine serum and 1% penicillin/streptomycin at 37 °C in 5% CO₂. For the stimulated condition, cells were treated with LPS (E. coli O111:B4, Sigma L4391) at 1 µg/mL for 16 h. The 0 h condition consisted of untreated cells collected immediately before LPS exposure.

### Mouse acute lung injury model

All procedures were approved by UAB IACUC and conducted in accordance with applicable guidelines. Male and female C57BL/6J mice (10-12 weeks old; Jackson Laboratory) were randomly assigned by cage to LPS or saline groups (n=5 per group). Sample processing and image acquisition were performed blinded to group assignment; group labels were unmasked for ML training. Under isoflurane anesthesia (2-3% in O2), mice received a single intratracheal instillation of LPS (E. coli O111:B4, Sigma L4391) at 1.5 mg/kg or an equivalent volume of sterile saline. At 24 h, peripheral blood was collected for monocyte staining and imaging flow cytometry, and serum was collected separately. Serum IL-6 and TNF-α were measured by DuoSet ELISA according to the manufacturer’s protocol. Mice were euthanized by CO2 inhalation followed by cervical dislocation. A designated lung lobe was weighed wet, dried to constant mass, and reweighed.

### Immunofluorescence microscopy

THP-1 cells were immobilized on coverslips, fixed in 4% paraformaldehyde, permeabilized with 0.1% Triton X-100, blocked in 5% BSA, and stained with anti-GM130 overnight at 4 °C followed by Alexa Fluor 594 secondary antibody. Nuclei were counterstained with DAPI. Images were acquired on an ECHO fluorescence microscope at 60× magnification. Exposure settings were not held constant across conditions, which limits the comparability of intensity-dependent features in the field-level analysis.

### GM130 morphometry

Cells were excluded according to prespecified quality control criteria (boundary contact, overlapping cells, poor focus, saturation, or failed segmentation). Ten prespecified morphometric and intensity features were extracted for each qualifying cell and summarized as the field-level median, with the imaging field serving as the unit of analysis. The 16 h fields were drawn from a single coverslip preparation; the 0 h fields were pooled from three acquisition batches.

### Imaging flow cytometry

Mouse peripheral blood was processed on the day of collection following established protocols for GM130- based Golgi morphometry on the ImageStreamX platform ^14^. Red blood cells were lysed, and samples were stained for viability (Ghost Dye Red 780), surface markers (CD45-APC, CD11b-FITC, Ly6G-PE), then fixed, permeabilized with saponin, and stained with anti-GM130. THP-1 cells were prepared similarly without surface staining. Images were acquired on an ImageStreamX at 60× magnification and analyzed in IDEAS version 6.2 using a prespecified gating sequence that selected focused single cells, excluded non-viable events, and identified CD45⁺CD11b⁺Ly6G⁻ cells.

### Machine learning classification

Conventional classifiers were trained on 41 numerical features extracted by IDEAS software and matched to the images. Deep learning models were trained on GM130 images of individual cells. The CNN learned spatial image features to distinguish control from LPS-treated cells. ResNet-18, MobileNetV3-Small, and EfficientNet-B0 used ImageNet-pretrained weights as initialization^30–32^. The THP-1 dataset contained 4,000 balanced images. The mouse dataset contained 1,366 images. In a separate analysis, an SVM trained on ten IDEAS features was used to assess permutation importance, measured as the decrease in test balanced accuracy after individually shuffling each feature 20 times. Bars show the mean and error bars the standard deviation. For visualization, a supervised feature encoder converted each cell’s IDEAS measurements into a vector of contributions to the classifier’s prediction. LightGBM served as the encoder, assigning each feature a signed contribution to the cell’s predicted LPS log-odds. These contributions sum to that score after adding a constant baseline, which was excluded from the encoding. These contribution vectors were standardized using statistics from the training set and projected into two dimensions using UMAP ^33^ and t-SNE ^34^, as previously applied to imaging flow cytometry data ^35^. UMAP was fitted on the training cells. Validation and test cells were subsequently transformed. The t-SNE projections were fitted jointly to all cells for visualization.

### Grad-CAM visualization

Gradient-weighted class activation mapping ^36^ was applied to visualize the spatial regions contributing to each model’s predictions. The method weights the final convolutional feature maps by their influence on the class score, highlighting image regions that contribute positively to the prediction. The resulting heatmap was overlaid on the cell image. Within-domain Grad-CAMs were generated using models trained and evaluated on the same dataset. For cross-domain analysis, activation maps were generated for the LPS class trained on the source dataset and their partially and fully adapted variants to the same representative target cells. The maps qualitatively show image regions supporting the predicted class.

### Model evaluation and statistical analysis

Data were partitioned into fixed 80:10:10 training, validation, and test splits at the image level. The balanced THP-1 dataset was divided into 3,200 training, 400 validation, and 400 test images. The mouse dataset was divided into 1,094 training, 136 validation, and 136 test images. Within-domain deep learning models were selected using validation balanced accuracy, with ROC-AUC as a secondary criterion, and training stopped when validation performance no longer improved. Accuracy denotes balanced accuracy. ROC-AUC, weighted F1 score, sensitivity, and specificity were calculated as complementary measures. To quantify sampling variability, 95% bootstrap confidence intervals were computed from 2,000 stratified resamples of the held-out test set predictions for each within-domain model. For cross-domain evaluation, models trained on one dataset were evaluated on the held-out test set of the other dataset. Cross-domain accuracy was evaluated separately for the no, partial, and full adaptation strategies. Under no adaptation, the source-trained model was evaluated on the target dataset without retraining. For deep learning, partial adaptation updated only the classifier head, whereas full adaptation updated the entire network. For conventional machine learning, partial adaptation recalibrated the source model’s prediction scores using labelled target training cells. Full adaptation refitted the classifier from scratch on those cells, so the conventional full-adaptation models are target-only rather than transfer models. The intensity baseline used logistic regression on mean intensity for full adaptation. Both deep learning adaptation strategies started from the model trained on the source dataset and used labelled target training cells. Training accounted for unequal class sizes and retained the normalization used for source training. Decision thresholds and adapted models were selected using weighted F1 on the target validation set, with accuracy and ROC-AUC as secondary criteria for model selection. The fixed target test sets were used only for final evaluation, not for fitting or selection. Cross-domain accuracy intervals used 1,000 stratified bootstrap resamples. ROC-AUC intervals used 2,000. Cross-domain ROC curves were computed exclusively from predictions after full adaptation on the independent target test cells. Field-level GM130 features were compared between conditions using two-sided Wilcoxon rank-sum tests, with P values corrected for multiple comparisons across ten features using the Benjamini-Hochberg procedure. Mouse physiological measurements were compared using two-sided Mann-Whitney U tests.

### Data availability

The imaging flow cytometry datasets and associated feature tables generated during this study are available from the corresponding author upon reasonable request.

## Acknowledgements

The authors thank the UAB Comprehensive Flow Cytometry Core for expert technical assistance and support.

## Author contributions

X.Z. and M.F. contributed equally. X.Z. designed and performed all cell culture, animal, imaging flow cytometry and immunofluorescence experiments, prepared samples and acquired data. M.F. developed and implemented the machine learning pipeline, performed all computational analyses and generated figures. B.G. conceived and supervised the computational analysis. P.C. conceived and supervised the study, designed experiments, interpreted results and prepared the manuscript with input from all authors. All authors reviewed and approved the final manuscript.

## Competing interests

The authors declare no competing interests.

## Funding

This work was supported by University of Alabama at Birmingham IMPACT Fund and ReInvent Fund.

---

**Supplementary Table S1.**
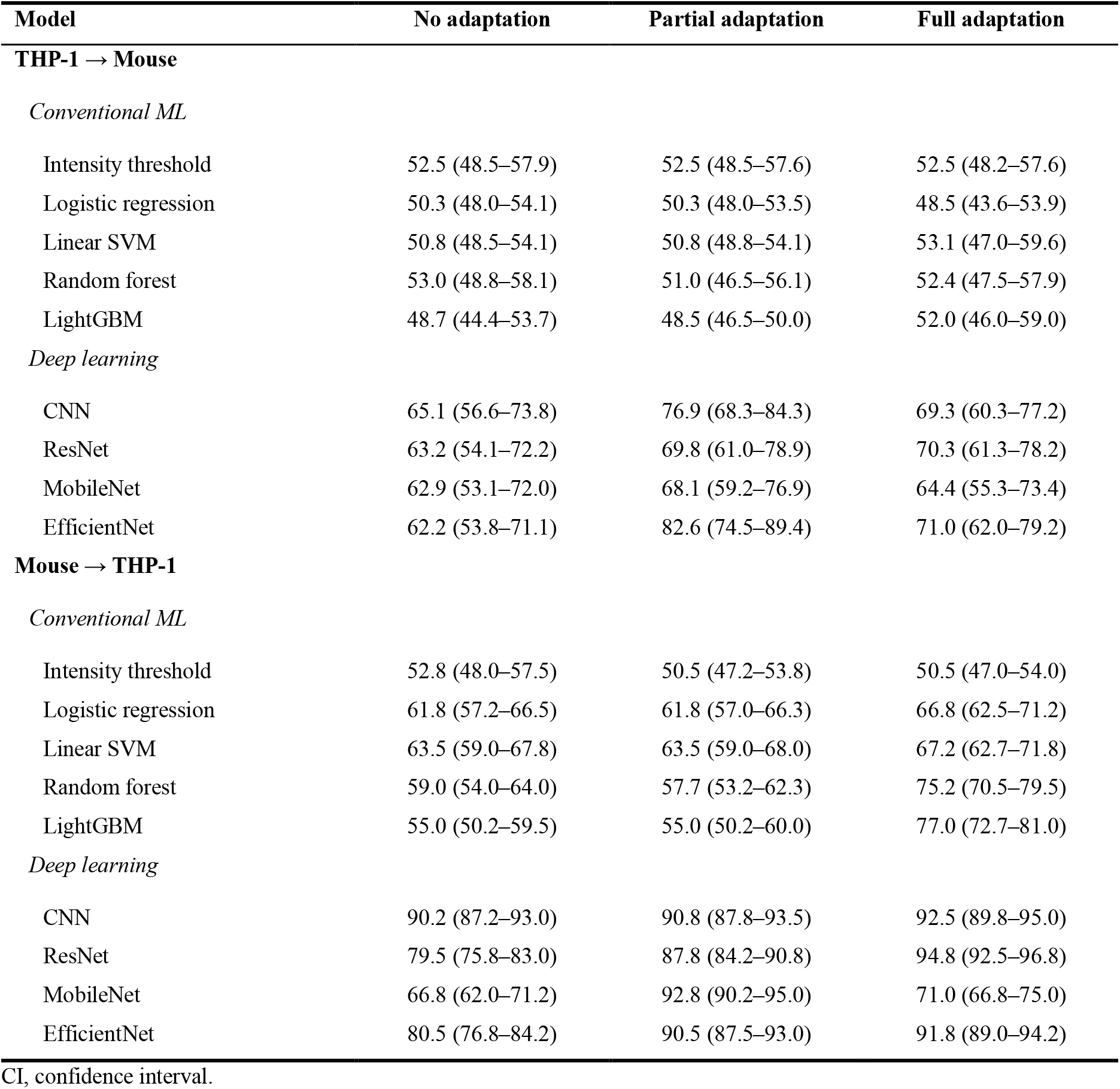
Cross-domain classification accuracy (%) with 95% confidence interval for models trained on one dataset and evaluated on the held-out test set of the other, under no, partial and full adaptation. No adaptation denotes fixed model parameters. Partial adaptation retrains the DL classifier head or calibrates ML scores; full adaptation retrains the full DL network or refits the ML classifier from scratch on target training data.

| Model | No adaptation | Partial adaptation | Full adaptation |
| --- | --- | --- | --- |
| **THP-1 → Mouse** |  |  |  |
| *Conventional ML* |  |  |  |
| Intensity threshold | 52.5 (48.5–57.9) | 52.5 (48.5–57.6) | 52.5 (48.2–57.6) |
| Logistic regression | 50.3 (48.0–54.1) | 50.3 (48.0–53.5) | 48.5 (43.6–53.9) |
| Linear SVM | 50.8 (48.5–54.1) | 50.8 (48.8–54.1) | 53.1 (47.0–59.6) |
| Random forest | 53.0 (48.8–58.1) | 51.0 (46.5–56.1) | 52.4 (47.5–57.9) |
| LightGBM | 48.7 (44.4–53.7) | 48.5 (46.5–50.0) | 52.0 (46.0–59.0) |
| *Deep learning* |  |  |  |
| CNN | 65.1 (56.6–73.8) | 76.9 (68.3–84.3) | 69.3 (60.3–77.2) |
| ResNet | 63.2 (54.1–72.2) | 69.8 (61.0–78.9) | 70.3 (61.3–78.2) |
| MobileNet | 62.9 (53.1–72.0) | 68.1 (59.2–76.9) | 64.4 (55.3–73.4) |
| EfficientNet | 62.2 (53.8–71.1) | 82.6 (74.5–89.4) | 71.0 (62.0–79.2) |
| **Mouse → THP-1** |  |  |  |
| *Conventional ML* |  |  |  |
| Intensity threshold | 52.8 (48.0–57.5) | 50.5 (47.2–53.8) | 50.5 (47.0–54.0) |
| Logistic regression | 61.8 (57.2–66.5) | 61.8 (57.0–66.3) | 66.8 (62.5–71.2) |
| Linear SVM | 63.5 (59.0–67.8) | 63.5 (59.0–68.0) | 67.2 (62.7–71.8) |
| Random forest | 59.0 (54.0–64.0) | 57.7 (53.2–62.3) | 75.2 (70.5–79.5) |
| LightGBM | 55.0 (50.2–59.5) | 55.0 (50.2–60.0) | 77.0 (72.7–81.0) |
| *Deep learning* |  |  |  |
| CNN | 90.2 (87.2–93.0) | 90.8 (87.8–93.5) | 92.5 (89.8–95.0) |
| ResNet | 79.5 (75.8–83.0) | 87.8 (84.2–90.8) | 94.8 (92.5–96.8) |
| MobileNet | 66.8 (62.0–71.2) | 92.8 (90.2–95.0) | 71.0 (66.8–75.0) |
| EfficientNet | 80.5 (76.8–84.2) | 90.5 (87.5–93.0) | 91.8 (89.0–94.2) |

CI, confidence interval.

## Notes

### Competing Interest Statement

The authors have declared no competing interest.

